# MEMO: A Micro Memo Sensor Detecting microRNA-RISC using an Accurate Cell-Free Expression Platform

**DOI:** 10.1101/2024.12.04.626909

**Authors:** Caroline E. Copeland, Chloe J. Heitmeier, Jeehye Kim, Yong-Chan Kwon

## Abstract

Circulating microRNAs have emerged as potential prognostic biomarkers of various diseases. They serve as micro memos sent out by cells, which scientists can intercept to gain real-time insights into cellular and disease status. However, establishing a standardized microRNA detection platform remains a challenge. Here, we integrate the RNA-induced silencing complex (RISC), which occupies mature microRNAs, into a complex cell-free synthetic genetic circuit using three different RNA polymerases, creating an innovative microRNA biosensor that is sensitive, precise, and cost-effective. The RISC protects the microRNA degradation during isolation and ensures the elimination of false target interactions, enhancing the detection robustness. With a limit of detection of 81 pM, affordability, and the possibility for on-demand use, this system proves to be a strong miRNA sensing tool.

## 1. Introduction

Monitoring and early detection of various illnesses are crucial medical practices to retain patients’ quality of health[1]. Addressing precision medicine gaps, soaring costs, and insufficient health insurance presents formidable obstacles to this pursuit[2]. New biosensing techniques for assessing health conditions using synthetic biology toolkits can alleviate the issues with current technologies and offer affordability owing to their shelf-stability and cell-based reactions with minimal to no equipment necessary to operate, as well as high specificity due to mimicking biomarker course of action[3]. Paired with an ideal circulating biomarker, microRNA, an accessible universal health monitoring tool has the potential to be created. MicroRNA (miRNA) is a type of small non-coding RNA that accounts for 1-3% of the mammalian genome[4] and was first discovered in 1993[5]. Recently, numerous studies have confirmed that the dysregulation of miRNA can be correlated with multiple cancer types, viral infections, diabetes, and cardiovascular diseases [6, 7, 8].

Many studies have shown that miRNA can be released into extracellular fluids, classifying them as circulating miRNA. They are found in various types of biological fluids[6] and are stable compared to other RNA molecules due to the protein complexes that encapsulate them. These miRNAs can resist degradation at room temperature for up to 4 days and survive relatively harsh conditions[7]. Additionally, many circulating miRNAs are found in extracellular vesicles (e.g., exosomes, apoptotic bodies, etc.), which provide an additional layer of protection and serve as a purification target[8]. The current knowledge about mRNAs’ significant role in gene regulation and high stability after cell export has positioned miRNAs as potential biomarkers for numerous diseases[9, 10]. These circulating miRNAs act as “micro memos” that scientists can detect to obtain real-time information on disease states.

Since their discovery, many approaches have been introduced to detect circulating miRNAs, but many suffer from performance consistency, high costs, or require well-equipped laboratory settings[11, 12]. New non-conventional methods have been developed to reduce cost or improve accessibility, but many still struggle with false positives and low specificity[13]. Specificity in miRNA detection is key to making a proper diagnosis or prognosis due to the high homology in the sequences of miRNAs[11]. The major reason for this lack of specificity arises from techniques relying solely on base-pairing of short, single-stranded, mature miRNAs. This hybridization method leads to problems where the probe only partly binds to a miRNA, making it difficult to determine if the signal is accurate, as well as problems of degradation by RNases[13, 14].

Previous miRNA sensing methods were overly synthetic and unfamiliar to the miRNA’s regular course of action. Mimicking the miRNA’s native functions would allow for the retention of accuracy and stability of the analyte. Hence, in this study, we develop a new miRNA detection method, a Micro Memo Sensor (MEMO), which includes using the entire RNA-induced silencing complex (RISC) holding the mature miRNA to perform its native functions in a cell-like reaction environment, the cell-free system (CFS), which not only protects the miRNA from degradation during isolation but also ensures the elimination of false target interactions, inducing an ultrasensitive gene circuit cascade in the reaction.

The CFS is a transcription-translation system using the machinery from lysed living cells. Since the cell is no longer viable, the user can direct the energy and cellular machinery solely toward the gene expression of interest by adding plasmid DNA[15]. With the system’s unprecedented level of freedom to modify and control biological systems, the CFS allows for the prototyping of complex cellular functions by breadboarding genetic parts[16-18], genetic circuits[19-22], protein modification[23-25], and biosynthetic pathways[26-29]. The CFS also allows for inexpensive sensing tests to be made due to the low-cost ingredients and accessibility with its ability to be freeze-dried and performed at room temperature[30]. Given these advantages and recent technical improvements, the CFS is an optimal tool for miRNA sensing and profiling.

The *Escherichia coli*-derived CFS miRNA sensing gene circuit presented here comprises a non-delaying, low-leak repression gene circuit that performs immediately in the OFF-state and can sensitively and promptly switch to the ON-state when the correct sequence miRNA-RISC is present. This sensing module circuit action is performed by a bacteria-based ECF11 sigma/anti-sigma (sigma11/anti-sigma11) repression system and its cognate[31]. In the ON state, gene transcription of a fluorescent protein, mNeonGreen (mNG)[32, 33], can be initiated when the circulating RISC-miRNA from the sample binds to a synthetically inserted complementary region on the anti-sigma mRNA after the start codon, causing the translation disruption of anti-sigma by either blocking ribosomes or slicing the sequence, allowing more sigma factors to be unrestrained and guide *E. coli* endogenous RNA Polymerase (endoRNAP) core enzyme to the DNA promoter ECF11 of mNG. This disruption by the RISC-miRNA is heavily sequence-dependent, allowing high fidelity that the miRNA triggering the circuit is the current sequence of interest. The signal is amplified by adding a third DNA construct containing the ECF11 sigma factor promoter, which triggers the expression of T7-CGG-split pseudo-sigma factor[34]. This pseudo-sigma factor couples with the core T7 split protein that is already supplemented to the reaction. After assembly, this complex can bind to its cognate CGG promoter and transcribe the mNG gene to produce a bright reporter protein (Figure 1a).

**Figure 1.**
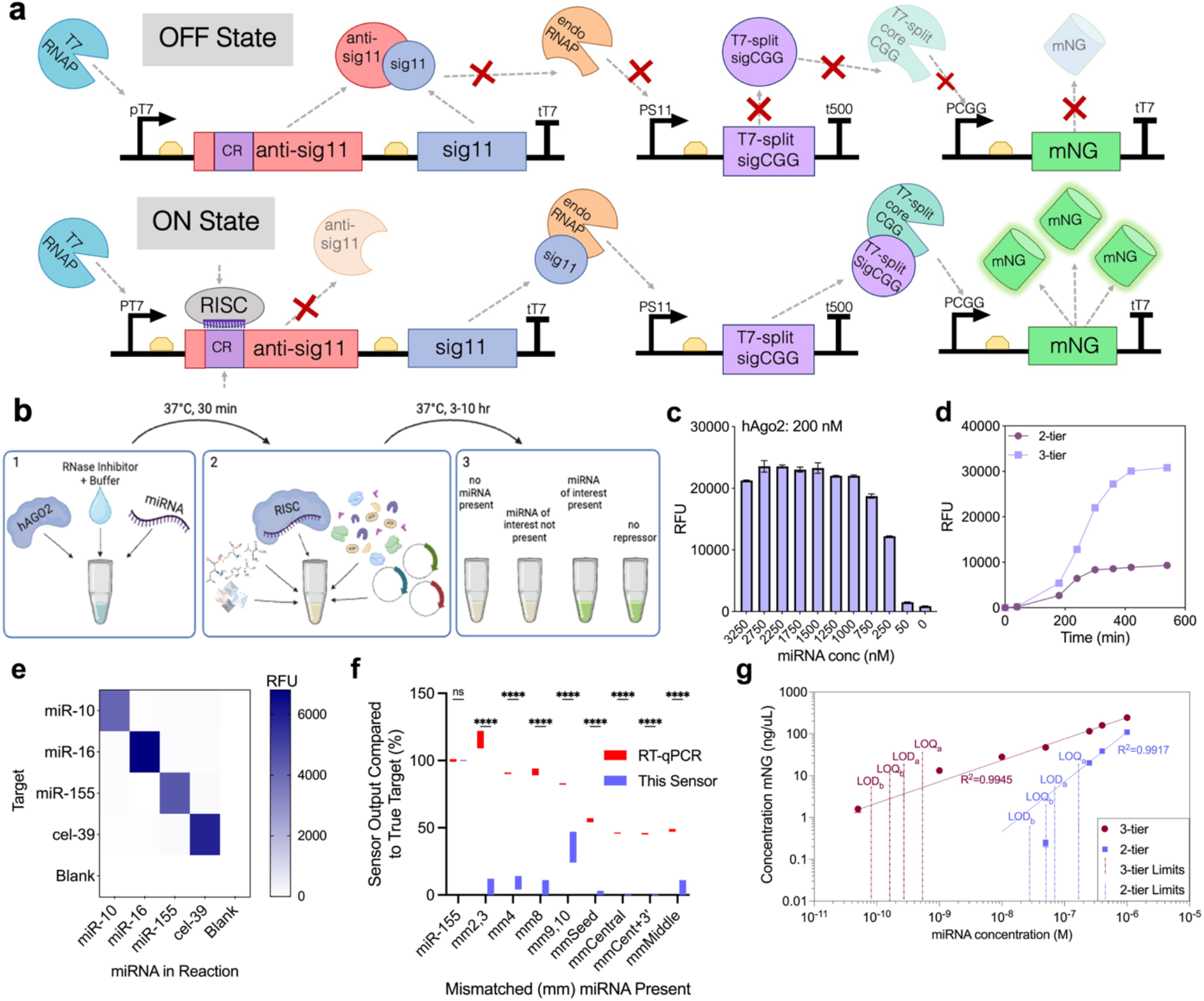
**a**: Schematic of the miRNA-RISC induced gene circuit.**b**: Diagram of miRNA sensing workflow. **c**: miRNA loading saturation experiment. **d**: Expression enhancements by 3-tier circuit. **e**: CFS miRNA sensor cross-reactions with varying miRNA targets. **f**: Accuracy of MEMO compared to RT-qPCR when differentiating between miRNA with similar sequences (mm=mismatch position). **g**: Limits of detection and quantification of the 3-tier circuit and 2-tier circuit. LOD and LOQ compared to blank “b” and compared to sensor activity in the presence of 1-nt different miRNA “a”. Values are represented as mean ± SD, n=3. ****p<0.0001.

## 2. Results and Discussion

### 2.1 MEMO Sensor Function and Characterization

The RISC comprises a mature miRNA serving as the guide (brains) and an Argonaute (AGO) protein providing the action (brawns). When bound to a target, miRNA can only form 1-2 helical turns, which does not result in much binding energy from nucleotide base stacking alone[35]. The AGO protein contains a lysine residue that neutralizes the negative charge of the bound miRNA at positions 1 and 3, allowing the guide to bind to the target without the negative charge repulsion of the backbones[36]. AGO also displays the seed nucleotides of the miRNA in a pre-helical structure, lowering the entropic barrier for target binding[37]. Thus, the binding to the target and turning of the miRNA is aided by the AGO protein coupling the miRNA.

A precise search and binding between RISC and the target and the coupling mechanism of AGO and miRNA facilitate the sensing accuracy of MEMO by only allowing the miRNA to bind to the target mRNA in sections. The sections include the 5’-end known as MID domain, seed section (beginning with sub-seed section), Mid or central, supplementary, tail, and PAZ domain[38]. The sub-seed nucleotides at the 2^nd^ through 4^th^ positions from the 5’-end practice lateral diffusion to find a target match[39]. Once the RISC finds the correct sub-seed match, it relieves a kink at the 7^th^ nucleotide, which allows the rest of the seed section to bind[39, 40]. If the seed match is not complimentary, the RISC complex releases the target and moves on to search for a new target in approximately one second[35, 41]. This mechanism ensures high specificity and low error rates. Once the entire seed sequence binds, the supplemental positions can bind and form a helix. The higher the target complementarity, the greater the binding becomes, which increases the time that the RISC complex remains on the target RNA[42] or induces a cleavage event of the target mRNA via AGO2 protein[37, 43].

To test the functionality of the cell-free MEMO sensor (Figure 1a), an artificial RISC was constructed by loading single-stranded miRNA into a purified human AGO2 (hAGO2) (Figure 1b)[41, 44]. The miRNA loading saturation point was found to be 1000 nM in the final cell-free reaction with 200 nM hAGO2 present (Figure 1c). The addition of an amplifier module after the sensing module, resulting in a 3-tier gene circuit (split T7 RNA polymerase, which will be discussed more in-depth in the following section), increases the output signal substantially while also increasing the reaction duration (Figure 1d). In addition, we confirmed the target specificity of MEMO by comparing cross-reaction among 4 different synthetic miRNAs (50 nM) (Figure 1e).

Next, we compared the detection accuracy with the conventional RT-qPCR method, which is currently considered one of the gold standards in miRNA detection. However, this method comes with significant limitations, especially in terms of accuracy and variability. Biomarker detection platforms require that both the limit of detection (LOD) and the dynamic range of the sensor fall within clinically relevant thresholds due to the typically low concentrations of biomarkers found in liquid biopsies such as blood and serum. RT-qPCR struggles to meet these requirements, often resulting in reduced accuracy and increased variability.

We evaluated the sensitivity of the MEMO platform by testing various concentrations of the miRNA-RISC complex and comparing the results with RT-qPCR with sequence mismatches that were altered at various numbered positions starting from the 5’-end (mismatch, mm). Although the RT-qPCR method has a clinically relevant detection limit, it exhibits a significantly narrow dynamic range, failing to differentiate between miRNAs with similar sequences compared to the actual miRNA target at tested levels, showing 90-100% signal for mismatches in the seed, and 50% signal for large sections of mismatches (Figure 1f). MEMO was able to differentiate and provide high dynamic ranges for all mismatches tested. However, MEMO showed a 40% signal only with a miRNA with a mismatch at the 9+10 position (mm9,10) (Figure 1f). Many studies have found that full complementary of the miRNA and target, especially at central nucleotides 9-12, is required for target cleavage[37, 42, 45]. Once the central region is bound, it allows the scissile phosphate to move into the catalytic site and the cleave occurs at between the phosphodiester bond between the bound target mRNA’s 10th and 11th (t10, t11) nucleotides. However, the manner in which the RISC binds to a target leaves this region last to bind[42].

Researchers have also discovered that a mismatch in this area is preferred for high affinity binding of a full complementary miRNA to avoid an unfavorable energetic arrangement[37], the opening of the central cleft of Ago2[46]. However, miRNA of similar sequence except for this central region may not exist in nature, as nature may not want to regulate the wrong gene. In this case, we found no miRNA of similar sequence to miR-155 except for the 9-10 region in the miRBase (mirbase.org). However, if there were a high sequence similarity, these miRNA have been found co-localized in the genome, and both may be dysregulated due to the disease state[47]. Also, miRNA with similar sequences share more similar target genes and target pathways, meaning that detecting any of the similar sequenced miRNA can lead you to the a similar physiological conclusion[47].

This hypothesis is supported by evidence indicating the 3’ supplementary region after g16 is rarely conserved across over 2 billion years of evolution, which is consistent with its minimal impact on cleavage or binding in Argonautes from various organisms[45, 48]. In contrast, RT-qPCR displayed signals even with considerable alterations in the miRNA sequence, highlighting MEMO’s advantage in leveraging the RISC’s capability to separate miRNA during binding, hence enhancing the sensing specificity and accuracy.

Finally, we established the LOD and limit of quantification (LOQ) for the MEMO circuit, both with and without the amplifier module (3-tier and 2-tier configurations, respectively), to highlight the critical role of the amplifier in reducing both the LOD and the LOQ. We measured the LOD compared to the blank ‘b,’ which was found to be 81 pM, and the LOQ was 271 pM. The equation used for the LOD (equation (1)) was based on a signal-to-noise ratio of 3, and the LOQ (equation (2)) ratio was based on a ratio of 10[49, 50]. These LOD and LOQ levels are comparable to other non-conventional miRNA methods[51]. With these detection thresholds, MEMO is capable of identifying a target miRNA expressing at 0.0144% of the total miRNA concentration in a 10 mL serum sample, considering a total miRNA concentration of 7 ng/mL, as previously determined[52] and detecting the target with quantification if expressing at 0.0475% (Figure 1g).

### 2.2. Sigma/Anti-sigma Factor Sensing Module Function in the Cell-Free System

The MEMO sensing module comprises a sigma factor/anti-sigma factor repression activation system, the first application of a sensing technique in the cell-free system. Many cell-free genetic circuit cascades have used a prokaryotic transcription factor repressor/inducer system, which typically requires the system to start in the ON state. However, the transcription factor repressors are less favorable for applications that require an initial OFF state because the repressor protein must first be expressed, transit with Brownian motion to the target DNA, and stay bound to it, which can cause a delay in repression for several minutes, while in the interim the reporter DNA is transcribed and translated[53]. On the other hand, the capability to start in the OFF state significantly reduces false positives, enhances the dynamic range, and improves the circuit sensitivity for detecting low levels of miRNA.

Achieving a strong OFF state is facilitated through sigma and anti-sigma DNA promoter regulation, where transcription initiation requires releasing the sigma factor from a strong protein-protein interaction[54]. Sigma and anti-sigma factor-based regulation have been validated in whole-cell studies, proving its strong OFF state in genetic switches[54-56]. The repression from anti-sigma is exceedingly strong, reaching a complete sequestering of the signal at a 4:1 anti-sigma:sigma molar concentration ratio supplementation of plasmid DNA containing the respective genes into the cell-free system. This is an endpoint measurement at 20h, indicating no regulatory leak due to repression disruption occurred at any point throughout the reaction (Figure 2a).

**Figure 2.**
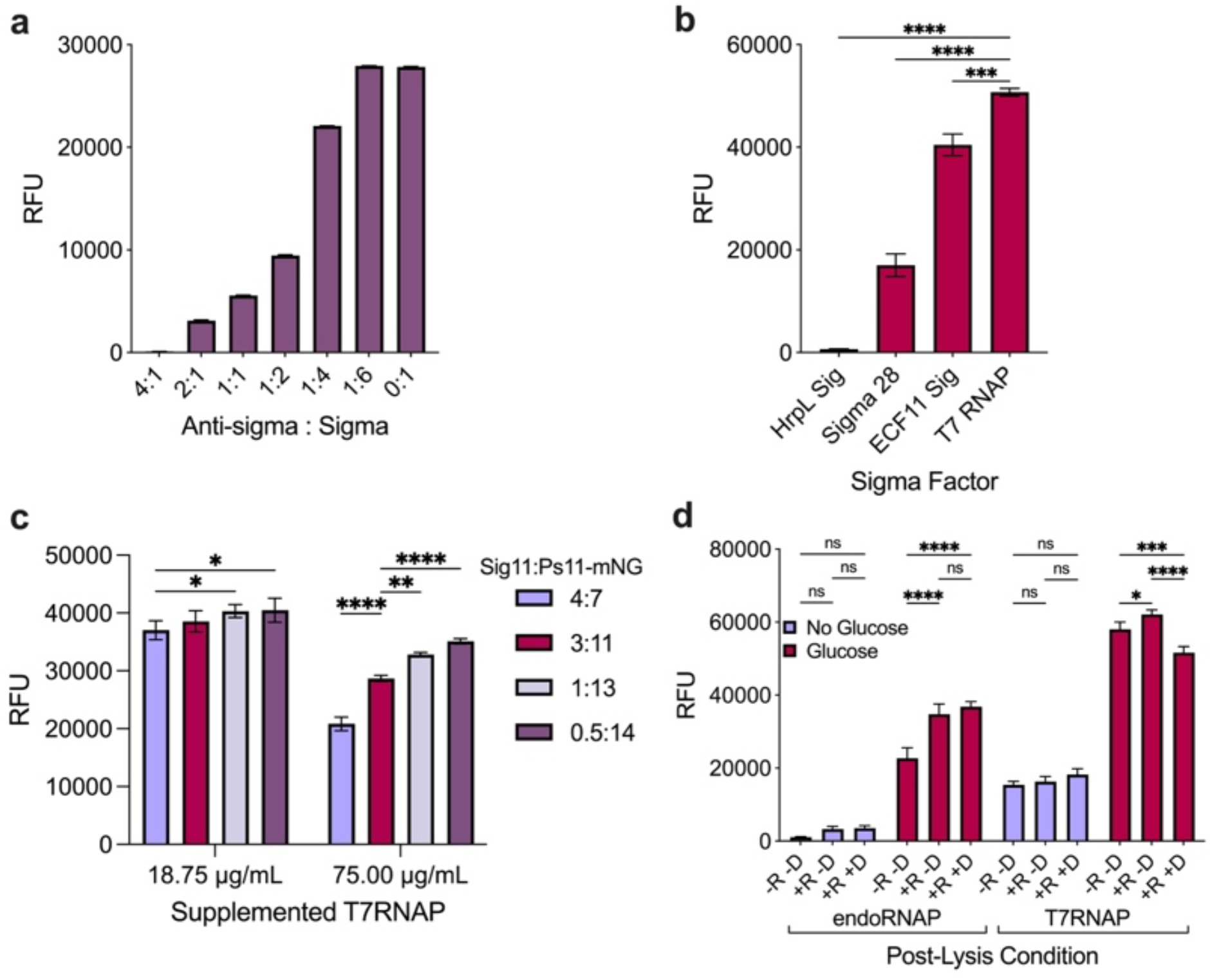
Characterizations of sigma/anti-sigma function in the CFS. **a**: Ratios of ECF11 sigma factor to anti-sigma factor in the CFS reaction expressing mNG in the active presence of sigma. **b**: Comparing sigma factor candidates’ activity in the CFS reaction to express mNG. **c**: mNG expression in the CFS with a dual RNAP system. ECF11 sigma factor was transcribed by two different concentrations of supplemented T7RNAP. Plasmid DNA concentrations of the T7-Sig11 plasmid and the Ps11-mNG plasmid were varied. **d**: Cell-free reaction performance of the system mainly derived by endoRNAP or T7RNAP using extracts made with glucose-rich or depleted media used during cell growth and varying post-lysis (R=runoff, D=dialysis). Values are represented as means ± SD, n=3, *p<0.05, **p<0.01, ***p<0.001, ****p<0.0001.

To demonstrate the robustness of sigma ECF11 in CFS, we compared two other sigma factors (*E. coli endogenous* sigma factor 28 and a non-native sigma factor HrpL from *Pseudomonas syringae*[57]) with T7 RNA polymerase (T7RNAP), which is commonly used in the CFS. All sigma factors were transcribed under the T7 promoter during the cell-free reaction. When the cognate promoter of each sigma factor was positioned upstream of the mNG fluorescent protein expression of mNG, sigma ECF11 demonstrated significantly higher expression levels compared to the other sigma factors (Figure 2b). Modeled in the 2-tier circuit without the amplifier, it is hypothesized that excessive mRNA production in the first step reduces the amount available for synthesis in the subsequent step, as shown in Figure 2c. It was found that when higher concentrations of T7RNAP and pJL1-Sigma11 plasmid were supplemented, lower expression of the second step (mNG expression from the pJL1-Ps11-mNG plasmid) was noticed. However, when we lowered the concentrations of supplemented T7RNAP, higher pJL1-Sigma11 concentrations were tolerated.

Next, we studied the cell extract preparation to achieve a robust protein synthesis output. Cell extract preparation varied from the standard process, especially when endoRNAP was the primary transcription engine in CFS, including the addition of post-lysis procedures such as runoff reaction and dialysis or the removal of glucose from the growth media[58, 59]. However, this system utilizes a combination of RNA Polymerases, so a varied characterization of optimal conditions was anticipated. It was found that glucose was crucial for maximizing protein synthesis output in both RNAP groups (Figure 2d).

### 2.3. Designing the Signal Amplifier Module: A 3-Tiered Circuit

To lower the detection limit of miRNA-RISC complexes, we integrated a signal amplification module following the initial sensing module using a second transcriptional activator type that can be activated by the first sigma11. Three potential candidates (RinA p80α[60] (RinA), T3 RNA Polymerase (T3 RNAP)[61], and split T7RNAP pseudo sigma factor CGG paired with the larger T7RNAP core piece[34]) were evaluated and chosen for exhibiting substantially lower crosstalk, ideally, no crosstalk between the endogenous *E. coli* components and the circuit designed in the CFS. The first candidate, RinA, a highly sensitive phage activator from *Staphylococcus aureus* phage 80α, had previously been utilized as a second amplifying module of a gene circuit in a whole-cell reaction[60] but did not perform effectively in the CFS. This diminished activity could be attributed to differences in system dynamics or the absence of co-factors that are typically present in whole-cell environments but were removed during post-lysis cell extract preparation[62]. Next, T3 RNAP was evaluated as an amplifying module, having previously been incorporated in cell-free gene circuit cascades[63, 64]. Unfortunately, T3 RNAP did not exhibit the strong response anticipated for the amplifying module, possibly due to high molecular weight, which consumed substantial system resources during transcription and translation. Finally, the Split-T7 RNA polymerase designed by Segall-Shapiro et al. was examined[34]. The system consisted of the split sections of the 285 amino acid DNA-binding loop, known as the ‘σ fragment,’ and the remaining 601 amino acid ‘core fragment’ with an attached SynZIP[34] for better joining of the two segments. Segall-Shapiro et al. also constructed σ fragments with distinct promoter specificities to remove crosstalk interactions with the regular T7 RNAP. The most effective sigma fragment, with the least crosstalk and highest performance, was the CGG sigma fragment, which has a CGG nucleotide-binding region within the regular T7 promoter. Recently, we have successfully implemented the SynZip system with a split fluorescent protein in the CFS, significantly enhancing the conjugation efficiency between the small and large protein segments[65].

The genes of interest were expressed using T7 RNAP within the CFS to evaluate the expression level of each candidate in comparison with the robust performance of sigma11. Each candidate gene was linked to its cognate promoter upstream of the mNG gene to facilitate comparing protein expression levels. As a result, the split T7 RNAP system was selected as the amplifier module due to its superior gene expression capabilities (Figure 3a).

**Figure 3.**
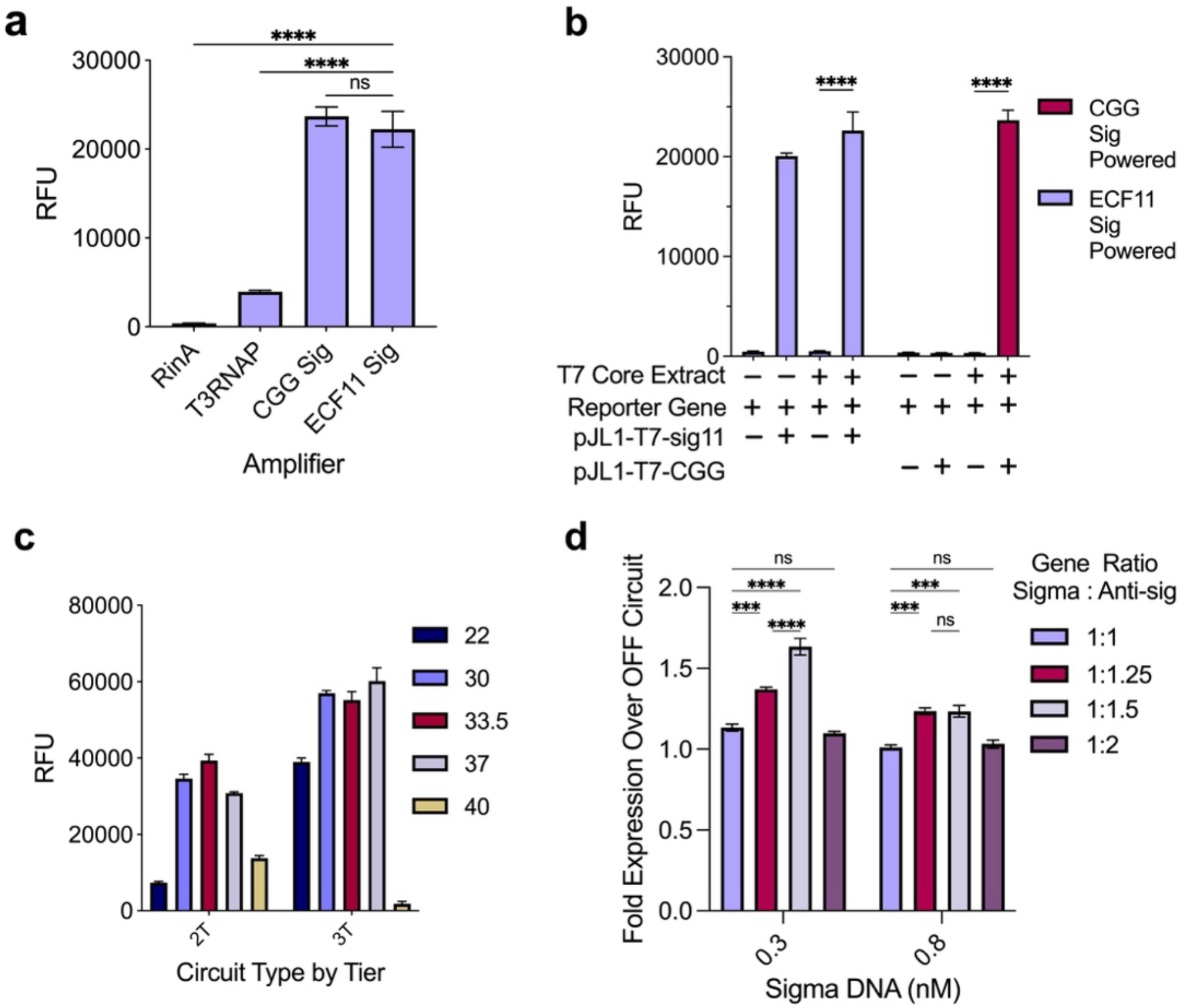
Characterizations of the 3-tiered amplifier module. **a**: Amplifier candidates’ activity in the CFS reaction compared to the initial activator ECF11 sigma factor with cognate promoters controlling mNG expression. **b**: CFS reaction testing T7RNAP split system. **c**: Circuit function by tier in the CFS expressing mNG reporter protein at various temperatures. **d**: Varying ratios of anti-sigma to sigma gene in the 3-tier circuit. Graphed data represents sensor fluorescent output in the presence of 50 nM of miRNA divided by sensor fluorescent output in the presence of 50 nM of miRNA differing by 1-nt at the 8^th^ position. Values are represented as means ± SD, n=3, *p<0.05, **p<0.01, ***p<0.001, ****p<0.0001.

To utilize the split T7 RNAP system, the T7 RNAP core fragment was overexpressed and supplied to the cell-free reaction. Simultaneously, the CGG sigma fragment gene was expressed during the reaction, facilitating in situ assembly upon the fragment synthesis. The plasmid harboring the CGG cognate promoter upstream of the mNG gene was also incorporated into the reaction. We confirmed that the split system exhibits no crosstalk with full-length T7 RNAP and no inhibition of the initial 2-tier circuit with sigma11 (Figure 3b). Since the 3-tier gene circuit and its parts used in this study have not been previously implemented in the CFS, a comprehensive characterization was necessary to optimize the circuit performance. The 3-tier system demonstrated enhanced expression at room temperature compared to the 2-tier configuration, offering potential reductions in incubation equipment costs (Figure 3c). Finally, in the 3-tier circuit, the ratio sigma11:anti-sigma11 was optimized and found to be 1:1.5, respectively, with the pJL1-sigma11 DNA having a lower concentration of 0.3 nM. This combination was found to be sufficient for optimal expression above a reaction exposed to a miRNA differing by 1-nt at the 8^th^ position (Figure 3d).

## 3. Conclusion

The MEMO platform represents a novel approach in miRNA sensing using an *E. coli* CFS. The system enables the entire RISC to retain its native functions, which hold the mature miRNA, and induce an ultrasensitive gene circuit cascade in the CFS. Previous miRNA detection platforms often failed to leverage the full potential of the RISC, which not only protects the miRNA from degradation during isolation but also reduces off-target detection by taking advantage of its inherent functions. To harness the RISC’s full capabilities, a cell-like working environment, yet a manipulable platform, is necessary. The CFS provides a user-defined platform that supports the intrinsic advantages of the RISC for sensitive miRNA detection. MEMO incorporates an ultrasensitive repression/activation method using the pair of non-native sigma/anti-sigma factors for the sensing module, coupled with a split T7 RNAP for the output signal amplification module, enabling precise and robust signaling. The MEMO platform achieves competitive detection limits with a LOD at 81 pM and LOQ at 271 pM. Moreover, due to the inexpensive system assembly and the feasibility of shelf-stable freeze-dried storage setting, MEMO surpasses many conventional miRNA detection methods, such as RT-qPCR, Next Generation Sequencing, and microarrays, in areas such as accuracy, low-variability, portability, simplicity, and cost-effectiveness. Looking forward, the MEMO platform is poised to become a powerful tool for microRNA detection and disease monitoring, promising significant contributions to biomedical research and clinical diagnostics.

## 4. Materials and Methods

### 4.1. Plasmid and Strain Information

Genes were assembled into a pJL1 vector backbone using Gibson assembly methods. Plasmids with genes expressed with T7 RNA polymerase in the cell-free system contained a T7 promoter and terminator. Plasmids with genes expressed by a sigma factor and endogenous RNA polymerase contained the cognate promoter for the plasmid being tested and the t500 terminator. The 2-tier gene circuit comprised four plasmids: pJL1-T7-AntiSigmaECF11 expressing anti-sigma ECF11 (687-nt, 25 kDa)[31], pJL1-T7-SigmaECF11 expressing ECF11 Sigma gene (585-nt, 22 kDa)[31] and the reporter plasmid pJL1-Ps11-mNG expressing the mNG gene[66]. The pJL1-Anti-SigmaECF11 plasmid contained a complementary miRNA-RISC binding site a few nucleotides after the stop codon. This position was screened to ensure no stop codon was made and no frameshift of amino acids would occur. The plasmids used in the amplifier circuit comprised: pJL1-T7-CGGsig expressing the CGG sigma fragment (996-nt, 37 kDa)[34], pJL1-Ps11-CGGsig expressing the same CGG sigma fragment but with the sigma ECF11 promoter, pJL1-PCGG-mNG expressing mNG gene by the CGG promoter[34]. The T7-split core fragment plasmid was obtained from Addgene (pTHSSd_38, #59961).

The bacterium *Escherichia coli* BL21 Star™ (DE3) (Thermo Fisher Scientific, Waltham, MA, USA), genotype F-ompT hsdSB (rB-, mB-) galdcmrne131 (DE3), was used for the cell-extract in the cell-free protein synthesis reaction. T7 RNA polymerase expression was controlled by the lacUV5 promoter in the bacteria genome. This cell line is also deficient in the gene rne131 which codes for RNase E, and the proteases Ion and OmpT, to decrease mRNA and protein degradation.

### 4.2. Cell-Extract Preparation

The extract used in this study was made as previously described[67, 68]. Briefly, cells were grown in 2xYTPG media and grown to the exponential phase. The cells were then harvested and washed three times with S30 buffer and stored until lysis, as described previously[58]. The cells were lysed by sonication[67]. Post-lysis steps were continued following protocols described previously[58, 67]. Cell extract aliquots were then flash-frozen in liquid nitrogen and stored at - 80°C. Total protein concentration was measured by Bradford Assay.

### 4.3. Split-T7 Core Expression and Purification

The T7 core split piece gene from Addgene plasmid pTHSSd_38, #59961, uploaded by Segall-Shapiro et al.[34] was inserted into a pETBlue1 vector with a N-terminus histidine tag after the start codon using Gibson Assembly. This plasmid was transformed into an IPTG-inducible BL21 Star™ (DE3). For protein purification, the cells were grown at 37°C with shaking at 300 rpm in a 2.5 mL Tunair shake flask. IPTG was added to the culture to induce T7 RNAP expression and subsequent expression of the T7 core split protein from the pETBlue1 plasmid. The next day, the cells were harvested, and His-tag purification was performed using Qiagen Ni-NTA resin per manufacture protocol. Once the sample was thawed, it was not re-frozen and saved to use again because the activity would be lost.

### 4.4. RISC Loading

Flag-tagged human Argonaute2 (hAGO2) protein was recombinantly made using a baculovirus expression system in Sf9 cells and given as a gift from the MacRae group[41]. These cells were then lysed with a Kontes Dounce tissue grinder and lysis buffer. Afterward, hAGO2 was purified using anti-flag M2 affinity gel and a competition FLAG peptide purchased as a kit from Sigma Aldrich. The eluted protein was then stored at 4°C and used within 2 weeks. Single-stranded guide microRNA-155 (5p) (ss-miRNA-155) with the 5’-end phosphorylated was purchased from Integrated DNA Technology. The loading reaction comprised RISC cleavage buffer (1 unit/µL RNasin, 20 mM Tris, 50 mM KCl, 5% glycerol, 1.5 mM MgCl2), 200nM Ago2, 1000nM of ss-miR-155, and nuclease-free water to make the final volume 2.15 µL. The mixture was then incubated at 37°C for 30 minutes for RISC formation.

### 4.5. Cell-Free Protein Synthesis

Reaction mixes were prepared following methods by Kim and Copeland (2019)[68], which included a salt mixture, NTPs, folinic acid, tRNA, HEPES, NAD, CoA, oxalic acid, putrescine, spermidine, PEP, 20 essential amino acids, T7 RNA polymerase, and 27% S30 cell-extract. The final volume was 15 µL. The reaction tubes were then incubated at 37°C for 20 h. For circuit reactions, the following were added: loaded RISC product at varying concentrations, 2.5 µg of purified T7 Split core piece 0.3 nM of pJL1-T7-SigmaECF11, 0.45 nM pJL1-T7-AntiSigmaECF11, 9 nM of pJL1-Ps11-CGGsig, and 14 nM pJL1-PCGG-mNG. OFF circuit reactions contained the same mixture as above, except for a miRNA that did not match the binding region by 1 nt at the 8^th^ position was added. Negative control for the circuit used all circuit plasmids without RISC formation product. Negative control for the reaction contained no DNA added. The negative control of the reporter protein expression was 13 nM of pJL1-Ps11-mNG and 2.15 µL of RISC formation product. As a positive control for the circuit, 2.15 µL of RISC formation product to a reaction without pJL1-T7-AntiSigmaECF11.

### 4.6. Protein Analysis

Once the reaction had reached 20 h, fluorescence was read with a SynergyTM HTX multi-mode microplate reader (BioTek, Winooski, VT, USA), with the filter having excitation and emission wavelengths at 485 nm and 528 nm, respectively. The remaining reaction mix was used to analyze total, soluble, and insoluble protein using SDS-PAGE (NuPAGE® 4–12% bis-tris gel (Thermo Fisher Scientific, Waltham, MA, USA)) to separate proteins in the mixture by molecular weight. A standard curve was also made for relative fluorescent units (RFU) as a function of protein concentration using a series of dilutions. Protein concentration was determined by overexpressing mNG in BL21 Star™ (DE3) culture, purifying with the Strep-tag attached to the protein, and finding the concentration using Bradford Assay.

### 4.7. LOD and LOQ Calculation

The limit of detection compared to the blank was found to be 81 pM, and the limit of quantification was 271 pM when following similar miRNA sensing paper guidelines for determining the LOD. However, this LOD and LOQ are based on the blank reaction (no miRNA), which is not a good representation of a true sample containing a variety of miRNA, including those similar in homology. To provide a better representative LOD and LOQ, the sensor response of detecting a miRNA with a 1-nt difference was considered the “blank” reaction. After these calculations, the LOD was determined to be 161 pM and LOQ 537 pM. The LOD equation (1) used was based on the signal/noise ratio of 3 and the LOQ equation (2) ratio of 10. This value was then interpolated using the x variables above and below the output y value LOD. In equations (1) and (2), µ_b_ is the mean of the blank, σ_b_ is the standard deviation of the blank, LOD is the limit of detection reporter value, and LOQ is the limit of quantification reporter value.

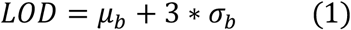

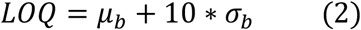

### 4.8. RT-qPCR and Efficiency/LOD Comparison Experiments

miRNA loading reactions were assembled and performed separately for each miRNA type, as described in section 4.4. This loading reaction was then treated as an initial sample to be added to the start of the sensing platform. This sample was immediately added to the reaction for cell-free reactions and incubated at 37°C. The miRNA was first purified from the AGO protein for RT-qPCR reactions as a liquid biopsy sample would normally be processed. A Qiagen miRNeasy Kit (cat. no. 217084) was used to purify each miRNA type. 1 µL (108 copies) of a control miRNA, UniSp6 RNA, was spiked in at the beginning of the purification to each tube to serve as a qPCR reference gene. After purification, the miRNAs were separately reverse transcribed into cDNA using the miRCURY LNA RT Kit (cat. no. 339340). Afterwards, the reaction was diluted at 1:30 for further qPCR reactions. miRCURY LNA miRNA PCR Assays (cat. no. 339306) and 2x miRURY SYBR® Green Master Mix (cat. no. 339345) were purchased from Qiagen (Germantown, MD, USA). For specificity tests, miRCURY LNA miR-155 primer from Qiagen was used for each miRNA reverse transcription reaction. UniSp6 RNA primers were also used to normalize the results to the spike-in. For testing the specificity of the miRCURY LNA miR-155 primers, ΔCt values were acquired from each reaction and compared to the value from the reaction with the true miR-155 sequence to output a percentage. Values were plotted in Figure 1f, representing the sensing units for both MEMO and RT-qPCR. The differentiation rate was calculated as a percentage by (1-(wrong target)/(correct target))*100.

### 4.9. Statistical Analysis

Statistical analyses were conducted using GraphPad Prism 8.4.3 (GraphPad Software) with a 5% significance level. For the parametric analysis of data from the quantification of the synthesized protein, two-way ANOVA followed by the Dunnett’s test was used.

## Author contribution

C.E.C. and Y.-C.K. conceived the project. C.E.C. and Y.-C.K. designed and conceptualized experiments. C.E.C. and C.J.H. prepared and performed experiments and acquired data. C.E.C. and J.K. analyzed and interpreted data. C.E.C. wrote the original manuscript. C.E.C. and Y.-C.K. revised and edited the manuscript. All authors contributed to the article and approved the submitted version.

## Acknowledgment

We would like to thank Ian Macrae and Luca Gebert for their generous supply of Sf-9 cell pellets with overexpressed and active hAGO2.

## References

1. Heitzer, E., S. Perakis, J.B. Geigl, and M.R. Speicher, The potential of liquid biopsies for the early detection of cancer, npj Precision Oncology, 1 (2017), p. 36. 10.1038/s41698-017-0039-5

2. Giljohann, D.A. and C.A. Mirkin, Drivers of biodiagnostic development, Nature, 462 (2009), p. 461–464.

3. McNerney, M.P., K.E. Doiron, T.L. Ng, T.Z. Chang, and P.A. Silver, Theranostic cells: emerging clinical applications of synthetic biology, Nature Reviews Genetics, 22 (2021), p. 730–746. 10.1038/s41576-021-00383-3

4. Bartel, D.P., MicroRNAs: genomics, biogenesis, mechanism, and function, Cell, 116 (2004), p. 281–97. 10.1016/s0092-8674(04)00045-5

5. Lee, R.C., R.L. Feinbaum, and V. Ambros, The C. elegans heterochronic gene lin-4 encodes small RNAs with antisense complementarity to lin-14, Cell, 75 (1993), p. 843–54.

6. O’Brien, J., H. Hayder, Y. Zayed, and C. Peng, Overview of MicroRNA Biogenesis, Mechanisms of Actions, and Circulation, Frontiers in endocrinology, 9 (2018), p. 402–402. 10.3389/fendo.2018.00402

7. Chen, X., Y. Ba, L. Ma, X. Cai, Y. Yin, K. Wang, J. Guo, Y. Zhang, J. Chen, X. Guo, Q. Li, X. Li, W. Wang, Y. Zhang, J. Wang, X. Jiang, Y. Xiang, C. Xu, P. Zheng, J. Zhang, R. Li, H. Zhang, X. Shang, T. Gong, G. Ning, J. Wang, K. Zen, J. Zhang, and C.Y. Zhang, Characterization of microRNAs in serum: a novel class of biomarkers for diagnosis of cancer and other diseases, Cell research, 18 (2008), p. 997–1006. 10.1038/cr.2008.282

8. Gallo, A., M. Tandon, I. Alevizos, and G.G. Illei, The majority of microRNAs detectable in serum and saliva is concentrated in exosomes, PloS one, 7 (2012), p. e30679. 10.1371/journal.pone.0030679

9. Wang, J., J. Chen, and S. Sen, MicroRNA as Biomarkers and Diagnostics, Journal of cellular physiology, 231 (2016), p. 25–30. 10.1002/jcp.25056

10. Witwer, K.W., Circulating MicroRNA Biomarker Studies: Pitfalls and Potential Solutions, Clinical Chemistry, 61 (2015), p. 56. 10.1373/clinchem.2014.221341

11. Detassis, S., M. Grasso, V. Del Vescovo, and M.A. Denti, microRNAs Make the Call in Cancer Personalized Medicine, Frontiers in Cell and Developmental Biology, 5 (2017), p. 10.3389/fcell.2017.00086

12. Pritchard, C.C., H.H. Cheng, and M. Tewari, MicroRNA profiling: approaches and considerations, Nature Reviews Genetics, 13 (2012), p. 358–369. 10.1038/nrg3198

13. Ouyang, T., Z. Liu, Z. Han, and Q. Ge, MicroRNA Detection Specificity: Recent Advances and Future Perspective, Analytical Chemistry, 91 (2019), p. 3179–3186. 10.1021/acs.analchem.8b05909

14. Reid, M.S., R.E. Paliwoda, H. Zhang, and X.C. Le, Reduction of Background Generated from Template-Template Hybridizations in the Exponential Amplification Reaction, Analytical Chemistry, 90 (2018), p. 11033–11039. 10.1021/acs.analchem.8b02788

15. Copeland, C.E., A. Langlois, J. Kim, and Y.-C. Kwon, The cell-free system: A new apparatus for affordable, sensitive, and portable healthcare, Biochemical Engineering Journal, 175 (2021), p. 108124.

16. Halleran, A.D. and R.M. Murray, Cell-Free and In Vivo Characterization of Lux, Las, and Rpa Quorum Activation Systems in E. coli, ACS Synth Biol, 7 (2018), p. 752–755. 10.1021/acssynbio.7b00376

17. Jayaraman, P., J.W. Yeoh, S. Jayaraman, A.Y. Teh, J. Zhang, and C.L. Poh, Cell-Free Optogenetic Gene Expression System, ACS Synth Biol, 7 (2018), p. 986–994. 10.1021/acssynbio.7b00422

18. Swank, Z., N. Laohakunakorn, and S.J. Maerkl, Cell-free gene-regulatory network engineering with synthetic transcription factors, Proc Natl Acad Sci U S A, 116 (2019), p. 5892–5901. 10.1073/pnas.1816591116

19. Siegal-Gaskins, D., Z.A. Tuza, J. Kim, V. Noireaux, and R.M. Murray, Gene circuit performance characterization and resource usage in a cell-free “breadboard”, ACS Synth Biol, 3 (2014), p. 416–25. 10.1021/sb400203p

20. Garamella, J., R. Marshall, M. Rustad, and V. Noireaux, The All E. coli TX-TL Toolbox 2.0: A Platform for Cell-Free Synthetic Biology, ACS Synth Biol, 5 (2016), p. 344–55. 10.1021/acssynbio.5b00296

21. Takahashi, M.K., J. Chappell, C.A. Hayes, Z.Z. Sun, J. Kim, V. Singhal, K.J. Spring, S. Al-Khabouri, C.P. Fall, V. Noireaux, R.M. Murray, and J.B. Lucks, Rapidly characterizing the fast dynamics of RNA genetic circuitry with cell-free transcription-translation (TX-TL) systems, ACS Synth Biol, 4 (2015), p. 503–15. 10.1021/sb400206c

22. Hu, C.Y., M.K. Takahashi, Y. Zhang, and J.B. Lucks, Engineering a Functional Small RNA Negative Autoregulation Network with Model-Guided Design, ACS Synth Biol, 7 (2018), p. 1507–1518. 10.1021/acssynbio.7b00440

23. Narumi, R., K. Masuda, T. Tomonaga, J. Adachi, H.R. Ueda, and Y. Shimizu, Cell-free synthesis of stable isotope-labeled internal standards for targeted quantitative proteomics, Synth Syst Biotechnol, 3 (2018), p. 97–104. 10.1016/j.synbio.2018.02.004

24. Oza, J.P., H.R. Aerni, N.L. Pirman, K.W. Barber, C.M. Ter Haar, S. Rogulina, M.B. Amrofell, F.J. Isaacs, J. Rinehart, and M.C. Jewett, Robust production of recombinant phosphoproteins using cell-free protein synthesis, Nat Commun, 6 (2015), p. 8168. 10.1038/ncomms9168

25. Kightlinger, W., K.E. Duncker, A. Ramesh, A.H. Thames, A. Natarajan, J.C. Stark, A. Yang, L. Lin, M. Mrksich, M.P. DeLisa, and M.C. Jewett, A cell-free biosynthesis platform for modular construction of protein glycosylation pathways, Nat Commun, 10 (2019), p. 5404. 10.1038/s41467-019-12024-9

26. Karim, A.S. and M.C. Jewett, A cell-free framework for rapid biosynthetic pathway prototyping and enzyme discovery, Metabolic engineering, 36 (2016), p. 116–126. 10.1016/j.ymben.2016.03.002

27. Karim, A.S., J.T. Heggestad, S.A. Crowe, and M.C. Jewett, Controlling cell-free metabolism through physiochemical perturbations, Metabolic engineering, 45 (2018), p. 86–94. 10.1016/j.ymben.2017.11.005

28. Dudley, Q.M., K.C. Anderson, and M.C. Jewett, Cell-Free Mixing of Escherichia coli Crude Extracts to Prototype and Rationally Engineer High-Titer Mevalonate Synthesis, ACS Synth Biol, 5 (2016), p. 1578–1588. 10.1021/acssynbio.6b00154

29. Casini, A., F.Y. Chang, R. Eluere, A.M. King, E.M. Young, Q.M. Dudley, A. Karim, K. Pratt, C. Bristol, A. Forget, A. Ghodasara, R. Warden-Rothman, R. Gan, A. Cristofaro, A.E. Borujeni, M.H. Ryu, J. Li, Y.C. Kwon, H. Wang, E. Tatsis, C. Rodriguez-Lopez, S. O’Connor, M.H. Medema, M.A. Fischbach, M.C. Jewett, C. Voigt, and D.B. Gordon, A Pressure Test to Make 10 Molecules in 90 Days: External Evaluation of Methods to Engineer Biology, J Am Chem Soc, 140 (2018), p. 4302–4316. 10.1021/jacs.7b13292

30. Silverman, A.D., A.S. Karim, and M.C. Jewett, Cell-free gene expression: an expanded repertoire of applications, Nature Reviews Genetics, 21 (2020), p. 151–170. 10.1038/s41576-019-0186-3

31. Rhodius, V.A., T.H. Segall-Shapiro, B.D. Sharon, A. Ghodasara, E. Orlova, H. Tabakh, D.H. Burkhardt, K. Clancy, T.C. Peterson, C.A. Gross, and C.A. Voigt, Design of orthogonal genetic switches based on a crosstalk map of σs, anti-σs, and promoters, Mol Syst Biol, 9 (2013), p. 702–702. 10.1038/msb.2013.58

32. Shaner, N.C., G.G. Lambert, A. Chammas, Y. Ni, P.J. Cranfill, M.A. Baird, B.R. Sell, J.R. Allen, R.N. Day, M. Israelsson, M.W. Davidson, and J. Wang, A bright monomeric green fluorescent protein derived from Branchiostoma lanceolatum, Nature Methods, 10 (2013), p. 407–409. 10.1038/nmeth.2413

33. Copeland, C.E., J. Kim, P.L. Copeland, C.J. Heitmeier, and Y.-C. Kwon, Characterizing a New Fluorescent Protein for a Low Limit of Detection Sensing in the Cell-Free System, ACS Synthetic Biology, 11 (2022), p. 2800–2810. 10.1021/acssynbio.2c00180

34. Segall-Shapiro, T.H., A.J. Meyer, A.D. Ellington, E.D. Sontag, and C.A. Voigt, A ‘resource allocator’ for transcription based on a highly fragmented T7 RNA polymerase, Mol Syst Biol, 10 (2014), p. 742.

35. Joo, C. and D. Rueda, Biophysics of RNA-Protein Interactions: A Mechanistic View, ed. C. Joo and D. Rueda. 2019: Springer-Verlag New York. VII, 249.

36. Nakanishi, K., Anatomy of RISC: how do small RNAs and chaperones activate Argonaute proteins?, Wiley Interdiscip Rev RNA, 7 (2016), p. 637–60. 10.1002/wrna.1356

37. Wee, L.M., C.F. Flores-Jasso, W.E. Salomon, and P.D. Zamore, Argonaute divides its RNA guide into domains with distinct functions and RNA-binding properties, Cell, 151 (2012), p. 1055–67. 10.1016/j.cell.2012.10.036

38. Klein, M., S.D. Chandradoss, M. Depken, and C. Joo, Why Argonaute is needed to make microRNA target search fast and reliable, Semin Cell Dev Biol, 65 (2017), p. 20–28. 10.1016/j.semcdb.2016.05.017

39. Chandradoss, S.D., N.T. Schirle, M. Szczepaniak, I.J. MacRae, and C. Joo, A Dynamic Search Process Underlies MicroRNA Targeting, Cell, 162 (2015), p. 96–107. 10.1016/j.cell.2015.06.032

40. Sun, G., J. Wang, Y. Huang, C.W. Yuan, K. Zhang, S. Hu, L. Chen, R.J. Lin, Y. Yen, and A.D. Riggs, Differences in silencing of mismatched targets by sliced versus diced siRNAs, Nucleic Acids Res, 46 (2018), p. 6806–6822. 10.1093/nar/gky287

41. De, N., L. Young, P.W. Lau, N.C. Meisner, D.V. Morrissey, and I.J. MacRae, Highly complementary target RNAs promote release of guide RNAs from human Argonaute2, Mol Cell, 50 (2013), p. 344–55. 10.1016/j.molcel.2013.04.001

42. Joo, C.R.D., Biophysics of RNA-protein interactions : a mechanistic view, (2019), p.

43. Dalmay, T., Mechanism of miRNA-mediated repression of mRNA translation, Essays Biochem, 54 (2013), p. 29–38. 10.1042/bse0540029

44. Yoda, M., T. Kawamata, Z. Paroo, X. Ye, S. Iwasaki, Q. Liu, and Y. Tomari, ATP-dependent human RISC assembly pathways, Nature Structural & Molecular Biology, 17 (2010), p. 17–23. 10.1038/nsmb.1733

45. Salomon, William E., Samson M. Jolly, Melissa J. Moore, Phillip D. Zamore, and V. Serebrov, Single-Molecule Imaging Reveals that Argonaute Reshapes the Binding Properties of Its Nucleic Acid Guides, Cell, 162 (2015), p. 84–95. 10.1016/j.cell.2015.06.029

46. Schirle, N.T., J. Sheu-Gruttadauria, and I.J. MacRae, Structural basis for microRNA targeting, Science, 346 (2014), p. 608-613. doi:10.1126/science.1258040

47. Kehl, T., C. Backes, F. Kern, T. Fehlmann, N. Ludwig, E. Meese, H.P. Lenhof, and A. Keller, About miRNAs, miRNA seeds, target genes and target pathways, Oncotarget, 8 (2017), p. 107167–107175. 10.18632/oncotarget.22363

48. Friedman, R.C., K.K.-H. Farh, C.B. Burge, and D.P. Bartel, Most mammalian mRNAs are conserved targets of microRNAs, Genome Research, 19 (2009), p. 92–105. 10.1101/gr.082701.108

49. Shrivastava, A. and V. Gupta, Methods for the determination of limit of detection and limit of quantitation of the analytical methods, Chronicles of Young Scientists, 2 (2011), p. 21.

50. McArdle, H., E.M. Jimenez-Mateos, R. Raoof, E. Carthy, D. Boyle, H. ElNaggar, N. Delanty, H. Hamer, M. Dogan, T. Huchtemann, P. Körtvelyessy, F. Rosenow, R.J. Forster, D.C. Henshall, and E. Spain, “TORNADO” – Theranostic One-Step RNA Detector; microfluidic disc for the direct detection of microRNA-134 in plasma and cerebrospinal fluid, Scientific Reports, 7 (2017), p. 1750. 10.1038/s41598-017-01947-2

51. Dave, V.P., T.A. Ngo, A.-K. Pernestig, D. Tilevik, K. Kant, T. Nguyen, A. Wolff, and D.D. Bang, MicroRNA amplification and detection technologies: opportunities and challenges for point of care diagnostics, Laboratory Investigation, 99 (2019), p. 452–469. 10.1038/s41374-018-0143-3

52. Mompeón, A., L. Ortega-Paz, X. Vidal-Gómez, T.J. Costa, D. Pérez-Cremades, S. Garcia-Blas, S. Brugaletta, J. Sanchis, M. Sabate, S. Novella, A.P. Dantas, and C. Hermenegildo, Disparate miRNA expression in serum and plasma of patients with acute myocardial infarction: a systematic and paired comparative analysis, Scientific Reports, 10 (2020), p. 5373. 10.1038/s41598-020-61507-z

53. Rosenfeld, N., M.B. Elowitz, and U. Alon, Negative autoregulation speeds the response times of transcription networks, J Mol Biol, 323 (2002), p. 785–93. 10.1016/s0022-2836(02)00994-4

54. Chen, D. and A.P. Arkin, Sequestration-based bistability enables tuning of the switching boundaries and design of a latch, Mol Syst Biol, 8 (2012), p. 620. 10.1038/msb.2012.52

55. Tiwari, A., G. Balazsi, M.L. Gennaro, and O.A. Igoshin, The interplay of multiple feedback loops with post-translational kinetics results in bistability of mycobacterial stress response, Phys Biol, 7 (2010), p. 036005. 10.1088/1478-3975/7/3/036005

56. Annunziata, F., A. Matyjaszkiewicz, G. Fiore, C.S. Grierson, L. Marucci, M. di Bernardo, and N.J. Savery, An Orthogonal Multi-input Integration System to Control Gene Expression in Escherichia coli, ACS Synth Biol, 6 (2017), p. 1816–1824. 10.1021/acssynbio.7b00109

57. Nissan, G., S. Manulis, D.M. Weinthal, G. Sessa, and I. Barash, Analysis of Promoters Recognized by HrpL, an Alternative σ-Factor Protein from Pantoea agglomerans pv. gypsophilae, Molecular Plant-Microbe Interactions®, 18 (2005), p. 634–643. 10.1094/mpmi-18-0634

58. Silverman, A.D., N. Kelley-Loughnane, J.B. Lucks, and M.C. Jewett, Deconstructing Cell-Free Extract Preparation for in Vitro Activation of Transcriptional Genetic Circuitry, ACS synthetic biology, 8 (2019), p. 403–414. 10.1021/acssynbio.8b00430

59. Shin, J. and V. Noireaux, Efficient cell-free expression with the endogenous E. Coli RNA polymerase and sigma factor 70, J Biol Eng, 4 (2010), p. 8. 10.1186/1754-1611-4-8

60. Wan, X., F. Volpetti, E. Petrova, C. French, S.J. Maerkl, and B. Wang, Cascaded amplifying circuits enable ultrasensitive cellular sensors for toxic metals, Nature Chemical Biology, 15 (2019), p. 540–548. 10.1038/s41589-019-0244-3

61. Garenne, D., S. Thompson, A. Brisson, A. Khakimzhan, and V. Noireaux, The all-E. coliTXTL toolbox 3.0: new capabilities of a cell-free synthetic biology platform, Synthetic Biology, 6 (2021), p. 10.1093/synbio/ysab017

62. Ferrer, M.D., N. Quiles-Puchalt, M.D. Harwich, M.Á. Tormo-Más, S. Campoy, J. Barbé, í. Lasa, R.P. Novick, G.E. Christie, and J.R. Penadés, RinA controls phage-mediated packaging and transfer of virulence genes in Gram-positive bacteria, Nucleic Acids Research, 39 (2011), p. 5866–5878. 10.1093/nar/gkr158

63. Shin, J. and V. Noireaux, An E. coli cell-free expression toolbox: application to synthetic gene circuits and artificial cells, ACS Synth Biol, 1 (2012), p. 29–41. 10.1021/sb200016s

64. Chizzolini, F., M. Forlin, N. Yeh Martín, G. Berloffa, D. Cecchi, and S.S. Mansy, Cell-Free Translation Is More Variable than Transcription, ACS Synthetic Biology, 6 (2017), p. 638–647. 10.1021/acssynbio.6b00250

65. Copeland, C.E., C.J. Heitmeier, K.D. Doan, S.C. Lee, K.B. Porche, and Y.-C. Kwon, Expanding the Cell-Free Reporter Protein Toolbox by Employing a Split mNeonGreen System to Reduce Protein Synthesis Workload, ACS Synthetic Biology, (2024), p. 10.1021/acssynbio.3c00752

66. Hostettler, L., L. Grundy, S. Kaser-Pebernard, C. Wicky, W.R. Schafer, and D.A. Glauser, The Bright Fluorescent Protein mNeonGreen Facilitates Protein Expression Analysis In Vivo, G3 (Bethesda), 7 (2017), p. 607–615. 10.1534/g3.116.038133

67. Kwon, Y.-C. and M.C. Jewett, High-throughput preparation methods of crude extract for robust cell-free protein synthesis, Scientific Reports, 5 (2015), p. 8663. 10.1038/srep08663

68. Kim, J., C.E. Copeland, S.R. Padumane, and Y.C. Kwon, A Crude Extract Preparation and Optimization from a Genomically Engineered Escherichia coli for the Cell-Free Protein Synthesis System: Practical Laboratory Guideline, Methods and protocols, 2 (2019), p. 10.3390/mps2030068

